# EpitoCore: mining conserved epitope vaccine candidates in the core proteome of multiple bacteria strains

**DOI:** 10.1101/864264

**Authors:** T.S. Fiuza, J.P.M.S. Lima, G.A. de Souza

## Abstract

In reverse vaccinology approaches, complete proteomes of bacteria are submitted to multiple computational prediction steps in order to filter proteins that are possible vaccine candidates. Most available tools perform such analysis only in a single strain, or a very limited number of strains. But the vast amount of genomic data had shown that most bacteria contain pangenomes, i.e. their genomic information contains core, conserved genes, and random accessory genes specific to each strain. Therefore, it is of the utmost importance to define core proteins, and also core epitopes, in reverse vaccinology methods. EpitoCore is a decision-tree pipeline developed to fulfill that need. It provides surfaceome prediction of proteins from related strains, defines clusters of core proteins within those, calculate the immunogenicity of such clusters, predicts epitopes for a given set of MHC alleles defined by the user, and then reports if epitopes are located extracellularly and if they are conserved among the core homologues. Pipeline performance is illustrated by mining peptide vaccine candidates in *Mycobacterium avium hominissuis* strains. From a total proteome of approximately 4,800 proteins per strain, EpitoCore mined 103 highly immunogenic core homologues located at cell surface, many of those related to virulence and drug resistance. Conserved epitopes identified among these homologues allows the users to define sets of peptides with potential to immunize the largest coverage of tested HLA alleles using peptide-based vaccines. Therefore, EpitoCore is able to provide automated identification of conserved epitopes in bacterial pangenomic datasets.

## INTRODUCTION

The characterization of specific molecular targets for controlling and removing bacterial infections is an important and challenging task. Membrane and surface proteins are known promising candidates due to the fact they carry out essential functions in the cell and are encoded by one-third of the bacterial genomes (Stevens and Arkin, 2000). The importance of those proteins is reinforced as they correspond to roughly 60% of known drug targets (Hopkins and Groom, 2002;Uhlen et al., 2015). Surface proteins are also key molecules for infection initiation and are at the interface with the host immune system (Finlay and McFadden, 2006). Furthermore, they are underrepresented in many experimental studies due to the fact that transmembrane proteins are heterogeneous, hydrophobic, and often detected at low abundance (Tan et al., 2008). Conventional screening of antigens in surface proteins is laborious, expensive and time-costly.

*In silico* approaches became a desirable method for mining candidate antigenic proteins. It has been largely employed to characterize single or sets of sequences of interest (Talukdar et al., 2014;Nagpal et al., 2018;Nosrati et al., 2019). Reverse vaccinology (RV) approaches use bacterial genomic information to achieve large scale antigen classification, and often integrate diverse levels of molecular prediction, such as subcellular localization, membrane adhesion and human cross-reactivity (for a review, see (Dalsass et al., 2019)). RV was first established to investigate antigens in serogroup B meningococcus (Pizza et al., 2000) and has been employed for many different pathogens (Seib et al., 2012;Delany et al., 2013).

Not surprisingly, many bioinformatics tools were developed in the past decade to facilitate epitope prediction in large datasets. They are either: i) decision-tree approaches, i.e., a large list of protein sequences from the organism of interest is submitted to the tool, which then apply different filters under specific parameters to trim the list to a final, smaller dataset of potential vaccine candidates; or ii) machine-learning approaches, which classify epitope candidates based on rules created by a training set of known, well characterized epitopes (Dalsass et al., 2019). Examples of such tools are NERVE (Vivona et al., 2006), VaxiJen (Doytchinova and Flower, 2007), Vaxign (He et al., 2010), Jenner-predict (Jaiswal et al., 2013) and many others. While some are based either in command line or web interface, most are often limited by the fact that they only allow the analysis of a single proteome of a bacterial strain. Vaxign allows multiple comparisons of strains in its web-server interface, however only a fraction of strains with complete sequenced genome are available for analysis (369 bacterial strains of 14,859 with complete and published genome as November 2019).

As more genome information of multiple strains of the same species was made available, the presence of a core and accessory genome was characterized in several of those species (McInerney et al., 2017). Only recently such genomic features are being taken into consideration when performing RV, as has been shown for *Helicobacter pylori* (Ali et al., 2015), *Acinetobacter baumanii* (Hassan et al., 2016), *Leptospira interrogans* (Zeng et al., 2017), pathogenic *Brucella spp*. (Hisham and Ashhab, 2018) and *Corynebacterium pseudotuberculosis* (Araujo et al., 2019). While these analyses provided the complete decision-tree performed, none of them provided the in-house scripts used for integration of all tools employed in their approaches. To our knowledge, the only tool available for pangenomic analysis and RV prediction is PanRV (Naz et al., 2019), which employs routinely used membrane and subcellular localization predictions. It also combines additional filters to enrich possible vaccine candidates such as gene essentiality and/or virulence factor predictions.

But PanRV (and other RV tools) are protein-centric, i.e. intact proteins are reported as vaccine candidates. If a vaccine is developed based on intact proteins, it could be argued that proteins containing more than two transmembrane domains are poor vaccine candidates due to the difficulty to purify them. However, synthetic peptides-based vaccines had been largely used in vaccine development in recent years. They offer many advantages to integral proteins purified from the pathogen, such as: i) fully in *vitro* manufacture, with less chance of biological contamination from pathogen; ii) full characterization as a chemical entity; iii) higher stability and storability; iv) smaller chance to induce non-specific reactions in the host (for a review, see (Skwarczynski and Toth, 2016)).

Through a peptide-centric analysis, even difficult to isolate proteins with highly immunogenic peptides could be considered for vaccine design. Based on this reasoning, we developed EpitoCore, a bioinformatic strategy that integrates surfaceome and subcellular localization prediction to pangenomic characterization, and further defines conserved epitopes in core proteins. For transmembrane proteins, EpitoCore correlates structure topology and epitope position to guarantee prediction of valid epitopes exposed to extracellular side.

## METHODS

### Scripts design and format

Scripts were created using Python version 3. CMG Biotools scripts are written in Perl and we added two in-house modifications to guarantee that: only the best alignment to a query sequence is reported instead of a list containing all alignments within the requested parameters; and that homologues are selected only if a bidirectional best hit criteria is fulfilled (see below). The TMHMM script was created using Perl as well. pSORTb is executed independently through a command line interface. IEDB peptide-HLA binding affinity predictors are written in Python 2.7. Final scripts for immunogenic analysis and image production were created using R studio version 1.1.442.

Users must provide a text file containing the species or the “intraspecies” name (as seen in the NCBI’s assembly summary information) to be investigated as input to the script get_proteome.py. This script outputs a comma-separated file with the assembly summary information requested and downloads the protein sequence datasets (.faa files) to a user’ specified folder. The .faa files are used independently for transmembrane and cell localization prediction. Script predict_transmembrane.py will call the TMHMM script, output the whole prediction in a folder, and filter proteins classified as transmembrane (see method below).Users must install and execute pSORTb, and script filter_psort.py will filter pSORTb outputs based on cutoff scores and localization given on parameters.

TMHMM transmembrane predicted proteins with a single helix are separated from the remaining TMHMM predictions by script filter_only_one_helix.py. Script intersect_psort_tmhmm.py compare TMHMM outputs with pSORTb and only keeps single helix proteins (SHPs) predicted with Unknown or Membrane localization. All scripts collect the sequences of those positively filtered proteins and saved them as a new .faa file in a separate folder.

Using the shortened .faa files, users must open the CMG Biotools suite to infer core proteins as described in detail below. All homologous proteins present in all strains (core proteins) will have their HLA binding affinity predicted by IEDB recommended tools using the immuno_prediction.py file. The immune_analysis.R script will combine the CMG Biotools core information with the IEDB antigenic information to discriminate protein clusters in which all homologues in all strains are highly immunogenic. It will also quantify epitope frequency per cluster (i.e. count of same peptide sequence present in proteins from a cluster) and epitope promiscuity (count of number of alleles recognized by same sequence). Comparison between epitope position and transmembrane topology can be optionally generated by the epitope_transmembrane_topology.R script.

### Data Acquisition

To compare full proteomes of *Mycobacterium avium hominissuis* strains, the amino acid sequences were obtained only for strains with complete genomes available (as November, 2018). Those are strains HP17 (GCA_002716905.1), OCU873s P7 4s (GCA_002716965.1) and OCU901s S2 2s (GCA_002716925.1) (Yano et al., 2017); H87 (GCA_001936215.1) (Zhao et al., 2017), TH135 (GCA_000829075.1) (Uchiya et al., 2013), MAC109 (GCA_003408535.1) (Matern et al., 2018) and OCU464 (GCA_001865635.2). Each protein dataset was retrieved from the National Center for Biotechnology Information (NCBI) database [21] using a python script that uses the Gene Assembly Summary file (assembly summary genbank.txt), available at ftp://ftp.ncbi.nlm.nih.gov/genomes/ASSEMBLY_REPORTS/. In total, each strain contained from 4,499 (OCU901s S2 2s) to 4,969 (HP17) annotated proteins.

### Identification of Transmembrane (TM) Domains

The protein datasets had alpha-helices transmembrane domains predicted by the standalone local variant of TMHMM version 2.0 (Sonnhammer et al., 1998;Krogh et al., 2001) (http://www.cbs.dtu.dk/services/TMHMM/). A second python script selected all sequences predicted to contain one or more TM alpha-helices as long as the helices comprised 18 or more amino acids. Predicted proteins were separated into two datasets: one with at least one helix occurred after the 60th N-terminal amino acid (fully embedded membrane proteins); and the other with only one helix prior to the 60th amino acid (helix close to possible true signal peptide) (SHPs). Such parameters are selected accordingly to TMHMM developers’ orientation. As TMHMM is a machine learning-based algorithm, such established parameters aim to reduce the number of false-positives as a result from expected pitfalls in the prediction. For example, short sequences rich in hydrophobic amino acids can be incorrectly classified as a protein containing transmembrane alpha-helices. Proteins with only one predicted helix close to protein N-terminal were further filtered as described below.

### pSORT analysis

All sequences were evaluated using the command line version of pSORTb version 3.0 (https://github.com/brinkmanlab/psortb_commandline_docker) (Nakai and Horton, 1999;Gardy et al., 2003). Gram-negative parameter setup was chosen, as it considered the recommended option for gram-positive bacteria with an outer membrane, such as *Mycobacterium spp*. The remaining parameters were kept as default. Proteins predicted to be located in periplasmic or outer membrane regions are kept. SHPs were also submitted to pSORTb independently, for localization control. Those predicted to be cytoplasmic are deleted, as well as those predicted to be periplasmic or outer membrane, since they are already included in the pSORTb dataset. SHPs predicted as cytoplasmic membrane or as unknown are kept.

### Definition of pangenomic components and surfaceome comparison

To better predict antigens present in all strains, we first define core and accessory proteins in all proteomes using the platform Comparative Microbial Genomics (CMG) Biotools (Vesth et al., 2013) version 2.2 (http://www.cbs.dtu.dk/biotools/CMGtools/). The Fasta files containing either complete proteome sequences or only predicted surfaceome entries were transferred to the platform, where CMG’ pancoreplot_createConfig script was executed, followed by CMGs’ pancoreplot script. CMG Biotools will consider two proteins as homologues when their BLAST alignment has at least 50% identity and 50% length coverage of the longest sequence. When BLAST aligns a protein from strain A with more than one protein in strain B, all proteins are considered homologues. We modified CMG’s pancoreplot script to report and cluster i) only the best aligned protein in strain B as an homologue; and ii) if same result is also true when strain B is used as query, i.e. a bi-directional approach where same result is achieved for strain A and B regardless which is used as query. All homologues are then clustered and classified as a single group. CMG also aligns a protein from a strain against all proteins from same strain, so sequences from within the same strain that fulfill the alignment threshold will be clustered together, meaning that some clusters may contain more than 7 proteins. CMG Biotools then outputs a group_*n*.dat file (where *n* is the cycle number) for every strain iteration, as well as a tbl file containing a summary and other intermediary documents. The data is cumulative for every iteration, therefore we use the last group_n.dat file to select clusters with proteins present in all strains. This analysis was performed for protein datasets previous to TMHMM prediction (whole proteome), or post TMHMM prediction.

### Immunogenetic Analysis

Immunological epitope prediction was carried out using recommended methods available at the Immune Epitope Database and Analysis Resource (IEDB) (Vita et al., 2019). It is well characterized that *Mycobacterium* species triggers MHC Class II CD4+ T cell responses in hosts, so we performed epitope prediction only to that MHC class. The IEDB recommended parameters for MHC-II uses the Consensus approach (Wang et al., 2008), combining NN-align, SMM-align, CombLib and Sturniolo. If no corresponding predictor is available for the allele, NetMHCIIpan is used (Andreatta et al., 2015). Different MHC/HLA alleles can be considered in this step. We selected 27 alleles for CD4+ T-cell epitope prediction, highly frequent in diverse populations and which were characterized as class II supertypes according to (Greenbaum et al., 2011). The alleles selected were DRB1*01:01, DRB1*03:01, DRB1*04:01, DRB1*04:05, DRB1*07:01, DRB1*08:02, DRB1*09:01, DRB1*11:01, DRB1*12:01, DRB1*13:02, DRB1*15:01, DRB3*01:01, DRB3*02:02, DRB4*01:01, DRB5*01:01, DQA1*05:01/DQB1*02:01, DQA1*05:01/DQB1*03:01, DQA1*03:01/DQB1*03:02, DQA1*04:01/DQB1*04:02, DQA1*01:01/DQB1*05:01, DQA1*01:02/DQB1*06:02, DPA1*02:01/DPB1*01:01, DPA1*01:03/DPB1*02:01, DPA1*01/DPB1*04:01, DPA1*03:01/DPB1*04:02, DPA1*02:01/DPB1*05:01, and DPA1*02:01/DPB1*14:01.

For each protein, the immunogenetic prediction profile lists all different epitope-MHC allele combinations with their respective affinity scores, as well as ranked immunogenetic percentiles. Here, we defined a protein’s immunogenetic Score as the mean scoring epitopes based on their percentile ranking (mean of all epitopes with score lower than 0.05). We did that for each protein within a cluster and then compared their Immunogenetic Scores - in a second filtering round, proteins whose scores were lower than 0.02 were then classified as Highly Immunogenic.

### Assigning epitope relevance to protein topology

Since the complete amino acid sequences of all predicted surfaceome proteins were submitted to epitope prediction, a vast number of highly immunogenic peptides were either present within the transmembrane region or in the intracellular portion of the molecule. Therefore we designed a script which aligns the protein topology prediction provided by TMHMM with the epitope prediction from IEDB. Epitopes which are located fully in the intracellular or transmembrane region, or at the interface of both, are excluded from the analysis. We only considered relevant epitopes if peptides were: fully aligned to the extracellular region; or, if partially embedded in the membrane, at least more than half of the peptide should be in the extracellular region (defined as parameter outside_ratio which should be higher than 0.5).

## RESULTS

### Protein sequences information and data analysis layout

Even though there were 201 genome entries for the species ‘*Mycobacterium avium’* in the Gene Assembly Summary of NCBI as of November 2018, we opted to perform antigenic analysis only for proteomes derived from strains with complete genomes sequenced. From 18 available datasets, seven belonged to strains of the subspecies *M. avium hominissuis* and were used as the raw input data in this work. The number of protein sequences available per strain ranged from 4,499 from strain OCU901s S2 2s to 4,969 from strain HP17 (Yano et al., 2017).

Figure 1 illustrates the data processing steps performed in this study. Briefly, each strain annotated proteome is submitted to either TMHMM prediction or to pSORTb prediction, to filter possible surfaceome from intracellular molecules. Surfaceome candidates are then compared across strains by CMG Biotools to define protein clusters containing homologues across strains. These proteins clusters are then classified accordingly to the number of proteins present in each cluster. We recommend surfaceome prediction to be run before clustering because, since CMG Biotools is run on a virtual machine (not locally), performance will be faster for smaller input files. Finally, initial epitome dataset are aligned to TMHMM topology prediction, and only extracellular sequences present in most or all homologues are considered valid epitopes.

**Figure 1.**
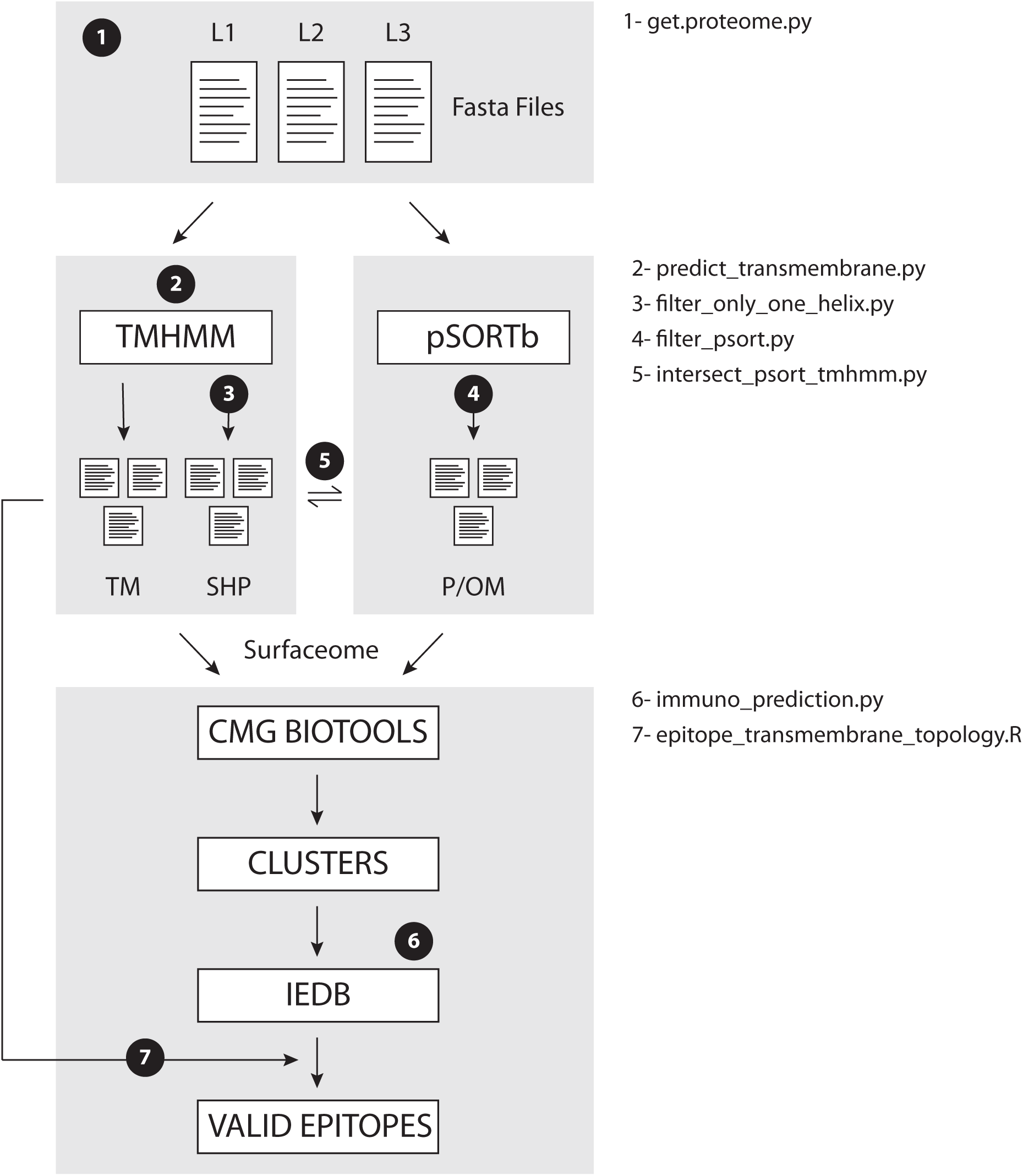
EpitoCore decision tree. Briefly, users retrieve fasta files for organism of interest, and perform transmembrane (TMHMM) and cellular localization (pSORTb) predictions separately. TMHMM output is further divided into proteins with helix outside or not of the signal sequence. Homologues within surfaceome predictions are clustered with CMG Biotools. Core clusters had their epitopes ranked by IEDB, and that output is aligned to protein topology. Conserved epitopes in all strains are then reported. Numbers 1-7 in workflow show location where each *in house* script is called.

### Performance of surfaceome and homology predictions

All protein entries present in the strains fasta files were submitted to transmembrane topology prediction and to cellular localization. Roughly 16% of each proteome was classified as sequences containing alpha-helix transmembrane domains or located to periplasmic and outer membrane regions (Supplementary Table I). For now we chose to exclude beta-barrel prediction from our approach, because such method is still hampered by the limited availability of known structural data (Tian et al., 2018), and because they are mostly observed in gram-negative bacteria rather than gram-positive bacteria such as *Mycobacterium avium* (Wimley, 2003;Freeman et al., 2011).

Surfaceome protein homologues were then clustered using CMG Biotools. Ideally, we wanted to characterize epitope presence in a cluster of homologues containing one protein per strain in all strains under investigation. For simplicity, from now on we illustrate in the figures the EpitoCore performance using the TMHMM prediction as an example. But all datasets generated in each step of the protocol are given in the Supplementary files available. Clustering of this specific dataset created a total of 577 groups, and most of those behaved as expected, i.e. the cluster was made of 7 proteins (397 clusters, Figure 2). Because CMG Biotools also BLAST a query entry against the remaining sequences from the same database, clusters containing more than 7 proteins were also observed. Only 10 clusters contained such possible paralogues. The remaining 170 clusters had six or less protein components and, in principle, is the accessory genome of the species. Supplementary File S1 lists all 577 clusters from the TMHMM prediction (in green), 294 clusters classified as SHPs (in orange), and 113 periplasmic and outer membrane clusters predicted from pSORTb (in purple). Accession numbers of the entries that belong to each cluster are also given.

**Figure 2.**
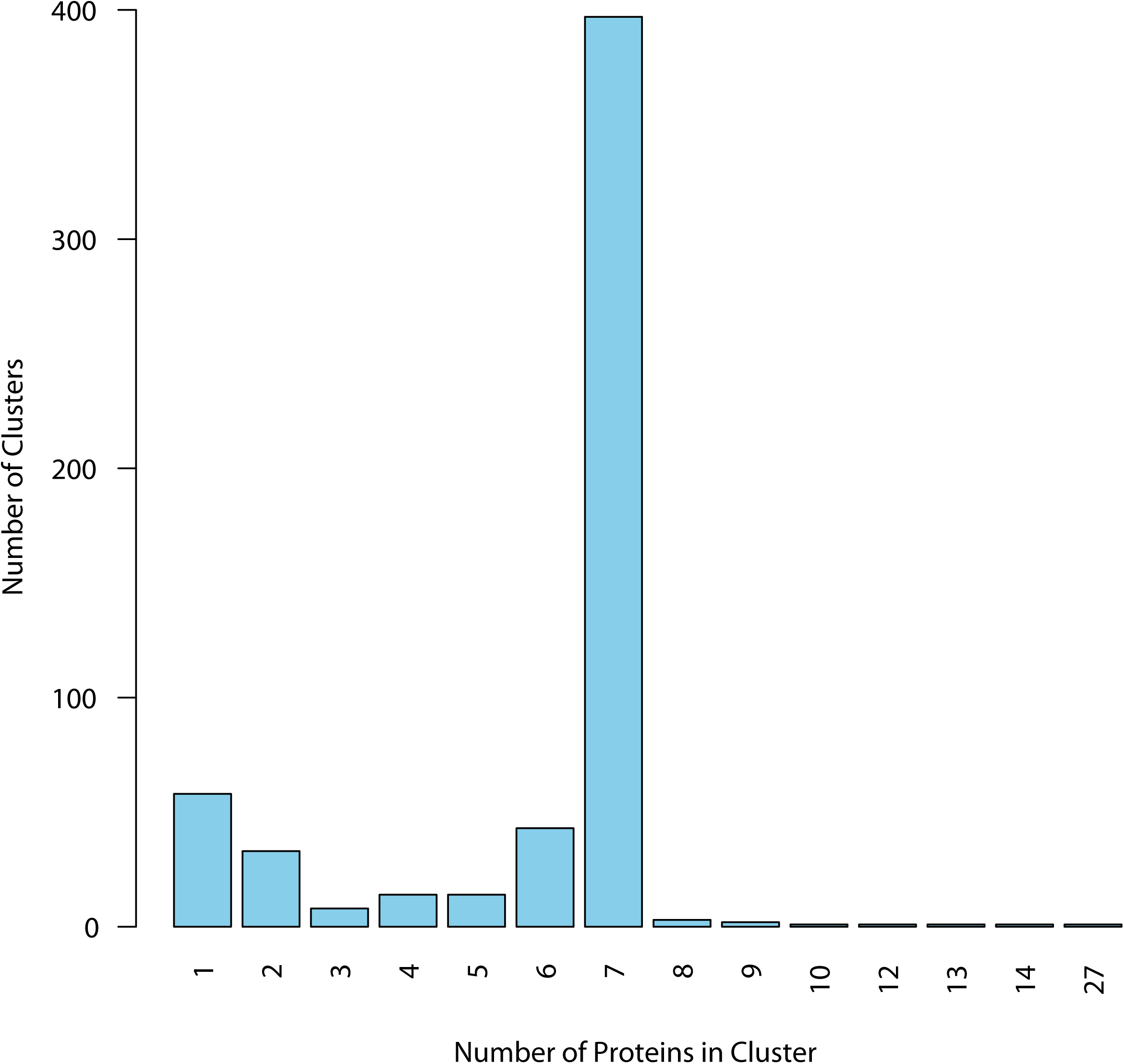
Distribution of homologues within clusters for TMHMM dataset. Proteins with membrane helix outside the signal peptide were clustered, and cluster composition was quantified. Majority of the clusters had, as expected, at least seven proteins, one from each of the analyzed strains, defining the core group. Clusters with six or less components are in principle defined as accessory proteins from the pangenome. Only core proteins were considered for immunogenic analysis.

Some of the accessory clusters (6 or less components) might be true core proteins and were mistakenly classified due to different reasons. We were able to detect at least two issues: i) annotation errors, meaning that the nucleotide sequence containing the gene exists in all strains, but was only annotated as a coding region in some of them; ii) group of homologues with conflicting prediction, i.e. some members containing transmembrane helices while the remaining did not, for example. This was common in clusters with lower sequence similarity between components. To demonstrate the last issue, we run CMG Biotools in the whole proteome dataset prior to surfaceome prediction, and compared cluster composition in both cases. Figure 3A shows that most clusters had the same composition regardless if clustered before or after the TMHMM protocol (zero missing proteins). The possible accessory clusters had observed patterns that could be explained by TMHMM variation (Figure 3B). See group 3, for example: all clusters with missing proteins had exactly four missing elements per cluster, meaning that when clustered before TMHMM, all clusters in group 3 had 7 elements. All proteins in that group are then core proteins, not accessory. A good number of such clusters could be re-classified as core due to different membrane domain prediction between homologues.

**Figure 3.**
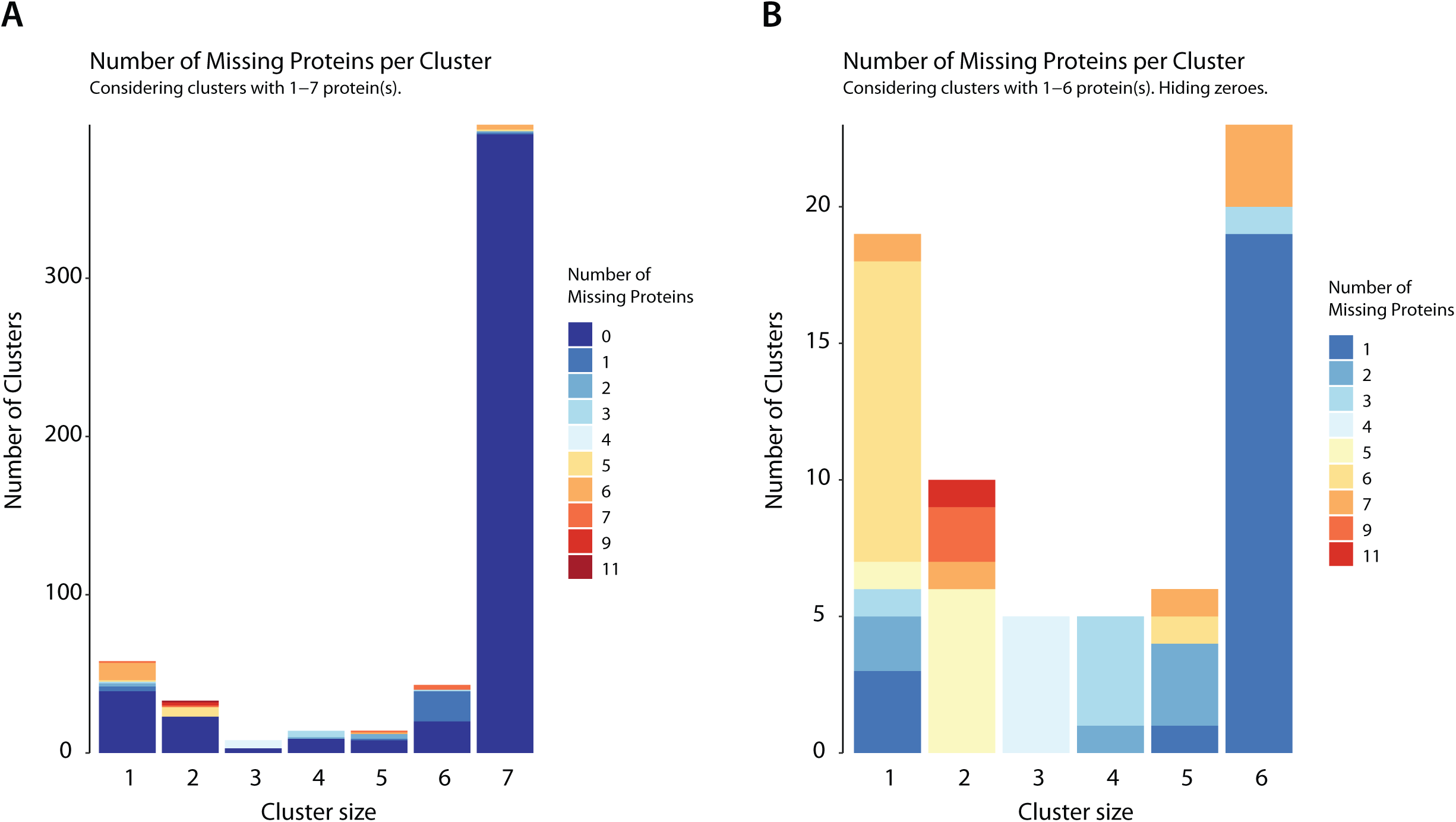
True core proteins with discrepant surfaceome prediction. To evaluate if clustering could be biased when performed on a smaller dataset, we also executed CMG Biotools in the whole proteome dataset of all strains. For the surfaceome proteins, cluster composition for most groups was identical regardless if prediction was done or not before clustering (zero missing proteins when clusters are compared) (A). This performance was close to 98% identical for core proteins. For accessory proteins (groups 1 to 6), this was closer to 55%. When hiding identical clusters (B), it is evident that for most accessory clusters, the number of missing elements adds to the exact number of strains used. This illustrates protein groups with 7 components if no prediction is performed, i.e. true core proteins.

### MHC-II epitope prediction

As seen in Figure 3A, Group 7 (i.e. clusters containing 7 protein sequences, one from each strain) was almost identical regardless if clustering is performed before or after any prediction. Only 6 clusters had missing proteins that were paralogues not classified as membrane proteins. For the aims of this work, epitope prediction was performed solely on clusters containing at least one homologue per strain. True core protein clusters where homologues had conflicting membrane domain or cellular localization prediction (i.e., groups 1 to 6 with missing proteins as exemplified in Figure 3B for TMHMM) were discarded.

All sequences present in clusters with +7 components were submitted to MHC-II epitope prediction using IEDB. For each cluster, its immunogenicity score was calculated by the median distribution of its peptides percentile ranking, to any given HLA allele, but only considering the top 5% better scoring peptides. Considering only better scoring peptides, most of the TMHMM predicted core clusters (387 of 407) would be considered “highly immunogenic” if a median score of 0.05 or less is used. We restricted the analysis to only a fraction of the clusters, therefore using a median score of 0.02 or less to define a highly immunogenic group. Such parameter selection defined that 112 transmembrane clusters were the most immunogenic of the dataset (Figure 4).

**Figure 4.**
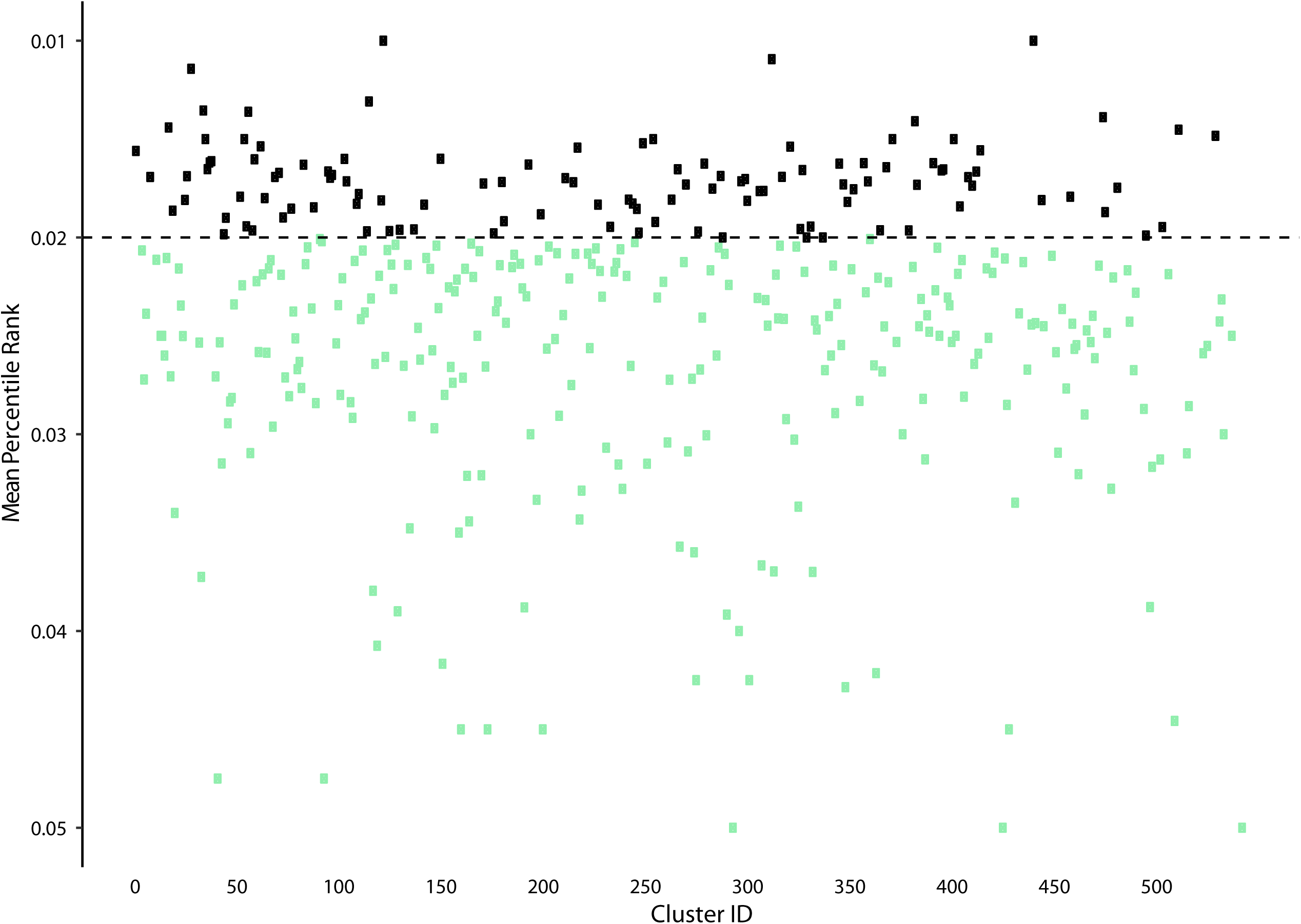
Immunogenicity Scores for TMHMM dataset. All sequences within each cluster were submitted to MHC II immunogenicity scoring, to all tested alleles. Sequences with percentile score lower than 0.05 were selected. Cluster immunogenicity is given by the mean distribution of all scored epitopes under this value, for all proteins. Highly immunogenic clusters were defined as those with a mean score lower than 0.02 (black dots).

### Epitope alignment to protein extracellular regions

Normally, when submitting a protein data for epitope prediction, the whole sequence is verified, including intracellular and transmembrane regions for transmembrane proteins. We wanted to characterize predicted epitopes which are accessible to the host immune system, i.e. located in the extracellular region of the protein. An additional filtering was then created in the approach, where amino acid locations provided by TMHMM topology prediction is aligned to each predicted epitope amino acid location in same protein.

An epitope was only considered as valid if at least more than half of its length is located extracellularly. When this parameter was applied to all highly immunogenetic clusters, approximately half of the clusters were deleted from further analysis because they contained no valid epitopes. Only 54 transmembrane, 34 SHPs and 15 periplasmic/outer membrane clusters remained, showing that most predicted epitopes are located in intracellular/transmembrane regions. Supplementary File S1 lists all the highly immunogenic clusters with valid epitopes. Supplementary File 2 lists all valid epitopes to each cluster component, which alleles they trigger, position in the protein, their outside score, frequency in cluster and promiscuity.

Supplementary Figure S1 illustrates a typical example, showing topology of epitopes for cluster 71 (cytochrome c oxidase subunit 3 family protein) before and after filtering. All seven proteins from this cluster had five predicted epitope windows, four of those located in transmembrane regions (“Before” panel). Only window 120-137 (18 amino acids containing 4 overlapping 15-amino acid epitopes) was located at the surface of the cell, and is the one that is considered a valid epitope after filtering (“After” panel). Such observations are not surprising, selective pressure should be higher for sequences at the interface with the host immune system, compared to intracellular and transmembrane regions. Therefore, observations of valid epitopes in extracellular portion of a transmembrane protein should be less frequent than in regions which are not under pressure of the host immune system.

### Epitope frequency and promiscuity in clusters

Overall, up to 75% of all considered epitopes are detected in at least 6 strains (Frequency column), while approximately 14% of the epitopes are unique to a single strain. Figure 5 shows epitope frequency in each of the transmembrane clusters containing valid epitopes. For 35 of those clusters, all predicted epitopes are conserved in the seven strains being evaluated. In addition, most clusters (43) contained at least one epitope conserved in all strains, and 48 clusters contained at least one epitope present in at least 6 strains. The remaining six clusters had epitopes which were predicted in no more than 5 strains, and those should be poor candidates for efficient immunization of all strains.

**Figure 5.**
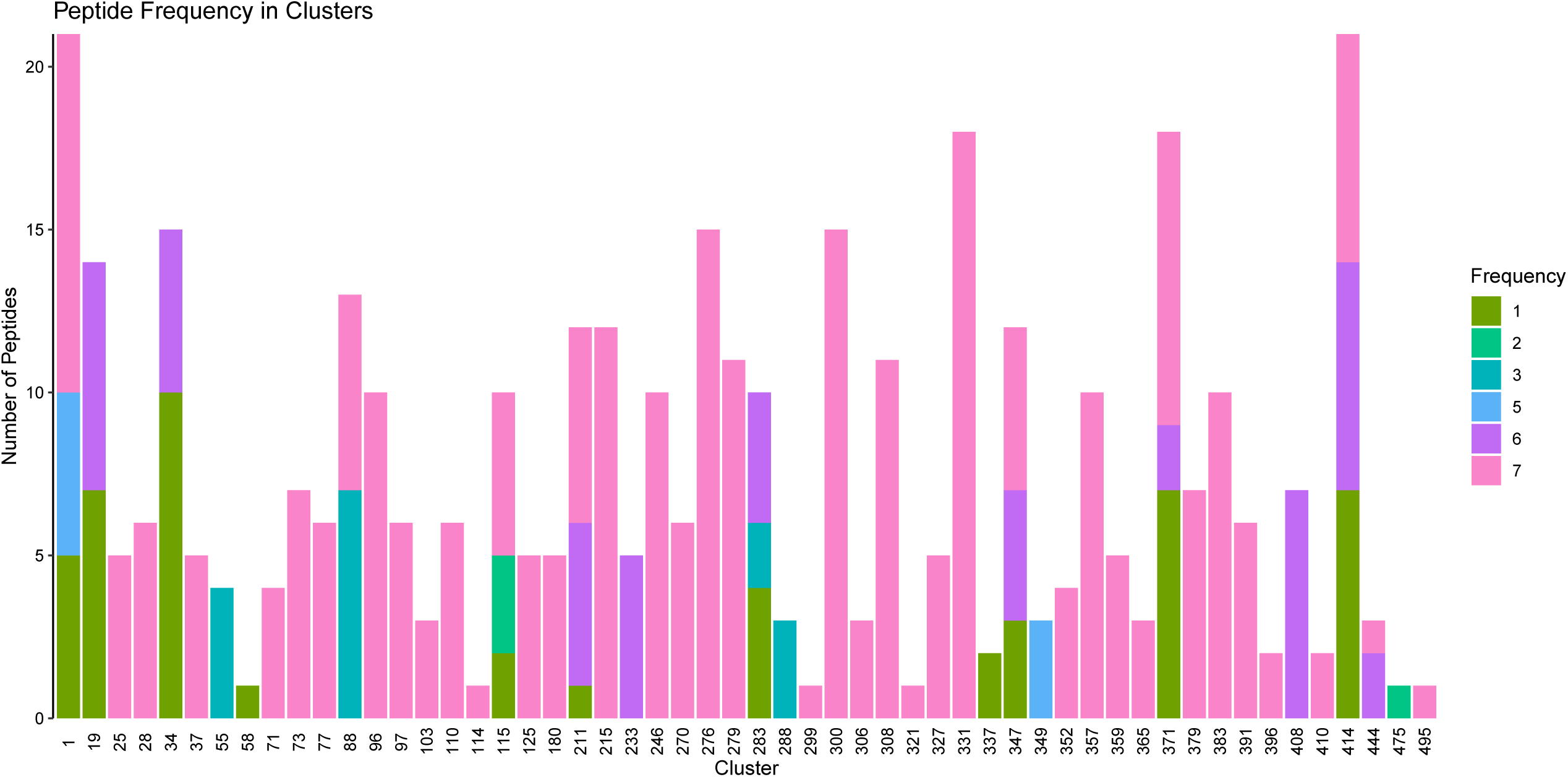
Valid epitope frequency within TMHMM clusters. Epitopes from highly immunogenic clusters were aligned to protein topology, and those intracellularly or inside the membrane were discarded. Remaining epitopes were then checked if they are conserved within all homologues of its cluster. Most clusters contained conserved identical epitopes in all homologues (count 7, pink). Epitopes present in 5 or less components of the cluster were not considered for further analysis.

However, epitope promiscuity, i.e. epitope recognized by more than one HLA allele, was observed for approximately 10% of the predicted sequences. No sequence could be an epitope for more than four alleles, and a single peptide (SVFRLFWLLYLGMTF) was an epitope to four HLA alleles (“HLA-DPA1*01:03/DPB1*02:01”, “HLA-DPA1*01/DPB1*04:01”, “HLA-DQA1*01:01/DQB1*05:01”, “HLA-DPA1*03:01/DPB1*04:02”). Promiscuous epitopes are present in 11 of the 54 clusters (Supplementary Figure S2), and interestingly seven of them are also clusters with exclusively conserved epitopes for all strains (Clusters 25, 71, 96, 180, 327, 391 and 410).

Finally, we also calculated the minimal set of epitopes potentially recognizable by the largest number of HLA alleles as possible. This calculation evaluates different combinations of epitopes with increasing group size until: a) all alleles tested are defined or; b) there is no gain in allele number as the number of epitopes being combined increases. With our valid epitopes dataset, we defined that 9 epitopes from 8 clusters are enough to immunize a population containing 15 of the 27 HLA alleles tested (Supplementary File 3). Manual checking of those epitopes using BLASTp showed that none shared sequence similarity to human proteins. This script will work accordingly to parameters inserted by the user at previous steps. For example, if larger population coverage is desirable, user can be less stringent when defining the highly immunogenic dataset. To achieve that, the mean IEDB immunogenicity score for the top 5% of predicted epitopes can be changed to 0.05 instead of the applied 0.02. In this case, 20 clusters containing valid epitopes would be enough to trigger immunological response from 22 of the 27 tested HLA alleles (data not shown).

## DISCUSSION

Searching for possible vaccine candidates using genomic data and bioinformatic pipelines is a well established approach (Dalsass et al., 2019). Nevertheless, the advances in next generation nucleotide sequencers had boosted the amount of available complete bacterial genomes. Consequently, it became evident that gene composition within genomes of strains of the same species can vary, characterizing pangenomes (McInerney et al., 2017). Until now, very few studies characterizing vaccine candidates had taken pangenomic features into consideration. The majority of bioinformatics pipelines were developed to investigate single genomes, or a very limited number of strains, and so far a single pipeline has been published for pangenomic RV analysis (Naz et al., 2019).

Therefore, we implemented EpitoCore to provide not only the prediction of antigenic proteins, but also to further mine conserved peptide vaccine candidates within core proteins of a species. We tested the pipeline using seven complete genomes of *Mycobacterium avium hominissuis* strains. Once a protein cluster (i.e. homologues from analyzed strains) is scored as immunogenic, we further restricted the pipeline to filter: epitopes that are distributed equally in all homologues; correctly aligned to protein topology; triggers MHC alleles that are representative in the population; and can be defined as a minimal set of epitopes for high population coverage immunization.

Initial steps in our decision-tree workflow followed routine standards in the field, by basically eliminating intracellular and inner membrane proteins by an alpha-helix transmembrane prediction using TMHMM (Sonnhammer et al., 1998;Krogh et al., 2001) and pSORT (Nakai and Horton, 1999;Gardy et al., 2003). We performed the surfaceome prediction prior to homology clustering in order to reduce dataset size and consequently, processing time. By performing homology clustering directly to the complete proteome of each strain, and comparing clusters with or without subcellular localization, we noted that many homologues had discrepant prediction due to sequence variations. While is not clear if this is taken into consideration by other publications using pangenomic features for RV, we recommend that not only core proteins to be considered as possible vaccine candidates, but core proteins with the same surfaceome prediction.

Core clusters were then submitted to MHC II epitope prediction using IEDB (Lundegaard et al., 2008;Vita et al., 2019). All protein homologues that are part of a cluster are challenged. Here, presence or absence of epitopes is not enough to simply define a cluster as immunogenic. Even when using a subset of MHC alleles frequent in the population (Greenbaum et al., 2011), practically all surfaceome proteins will have epitopes. EpitoCore lists all epitopes predicted by IEDB and sort those with lower percentile rankings (<0.05) for each protein within each cluster. The cluster immunogenicity is then given as the average percentile score of those epitopes for all proteins.

Here, in addition to define protein groups with valid epitopes, it is important to distinguish: a) if epitopes are in accordance to protein topology, i.e. is located extracellularly; and b) if epitopes are conserved amongst the homologues. For membrane proteins and SHPs, we observed that most epitopes were within transmembrane regions and discarded them. Only a fraction of clusters with epitopes predicted by IEDB could be considered to possess valid extracellular epitopes. Most epitopes are situated in the transmembrane or intracellular portions, which may be explained by an evolutionary pressure exerted on the bacteria by the host immune system (Halling-Brown et al., 2008). Some of these clusters are homologues to *Mycobacterium tuberculosis* proteins known to be related to: virulence, such as mammalian cell entry proteins (clusters 710, 714, 717, 734, 753, 791,798, 836, 842, 849 and 864) (Zhang and Xie, 2011) and type VII secretion proteins (clusters 595, 644, 773 and 774) (Ates et al., 2016); drug resistance (clusters 283, 578, 707, 722) (Hugonnet et al., 2016;Briffotaux et al., 2017); drug targets (clusters 691) (Gordon et al., 2000), to name a few.

The distribution of the remaining epitopes in the homologues is random, with some present in all strains while other present in just some of the related proteins. Ideally, for vaccine development, only epitopes present in most, if not all, strains should be considered. EpitoCore ranks epitopes based on their frequency, and only those conserved in the cluster are reported to users. Epitope promiscuity, rather than density, is also checked. Both features can be used to calculate a minimal number of epitopes which binds the largest number of MHC alleles tested. In our dataset, we determined that 9 epitopes are predicted to bind 15 of the 27 alleles tested.

## CONCLUSION

EpitoCore is a peptide-centric epitope prediction tool which takes into consideration pangenomic variation across strains of a given dataset, and reports conserved epitopes between homologues of those strains. The pipeline is highly modular, and future developments could allow implementation of transcriptomics/proteomics checks to verify that clusters containing predicted epitopes are indeed expressed in the organism of interest.

## AUTHORS CONTRIBUTION

TSF designed the pipeline and evaluated the data. JPLS assisted with project development and discussions. GAS designed the project, evaluated the data and assisted with project development and discussions.

## ACKNOWLEDGMENTS

This work was supported by grants 23038.004629/2014-19 to GAS and 88887.162254/2017-00 to TFS from the Coordenação de Aperfeiçoamento de Pessoal do Ensino Superior. GAS was also supported by grant 406630/2016-0 from Conselho Nacional de Desenvolvimento Científico e Tecnológico (CNPq).

## FIGURE LEGENDS

**Supplementary Figure S1 – Alignment of the protein topology to the epitope prediction**. An illustrative example of a homologue from cluster 71 (Cytochrome c subunit III protein). Immunogenicity prediction defined four regions of this protein to contain possible epitopes (Before panel, in pink). As shown in the figure, three of those regions are within the transmembrane domain. Protein topology alignment allowed the removal of such regions, which were then considered invalid epitopes (After panel).

**Supplementary Figure S2 – Epitope promiscuity within TMHMM cluster dataset**. Valid epitopes with frequencies higher than 6 were checked for binding capacity of more than one MHC allele. Majority of epitopes were predicted to bind a single allele (red), with only a few epitopes from 11 clusters predicted to bind two or three alleles. Only a single epitope from cluster 327 was predicted to bind four alleles.

