## Supplementary figures and images for "EpitoCore: mining conserved epitope vaccine candidates in the core proteome of multiple bacteria strains"

### Supplementary Figure S1

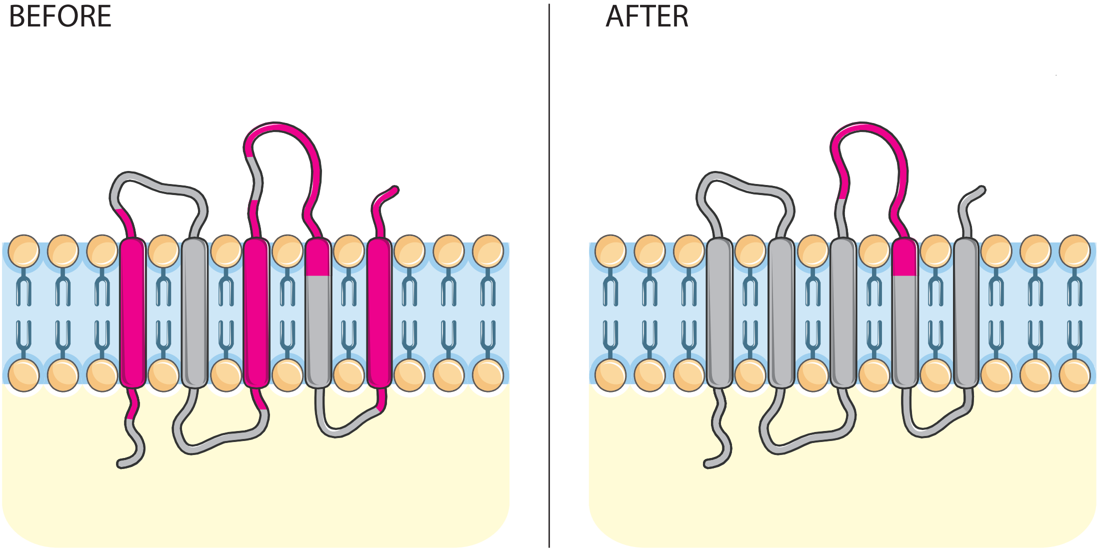

### Supplementary Figure S2

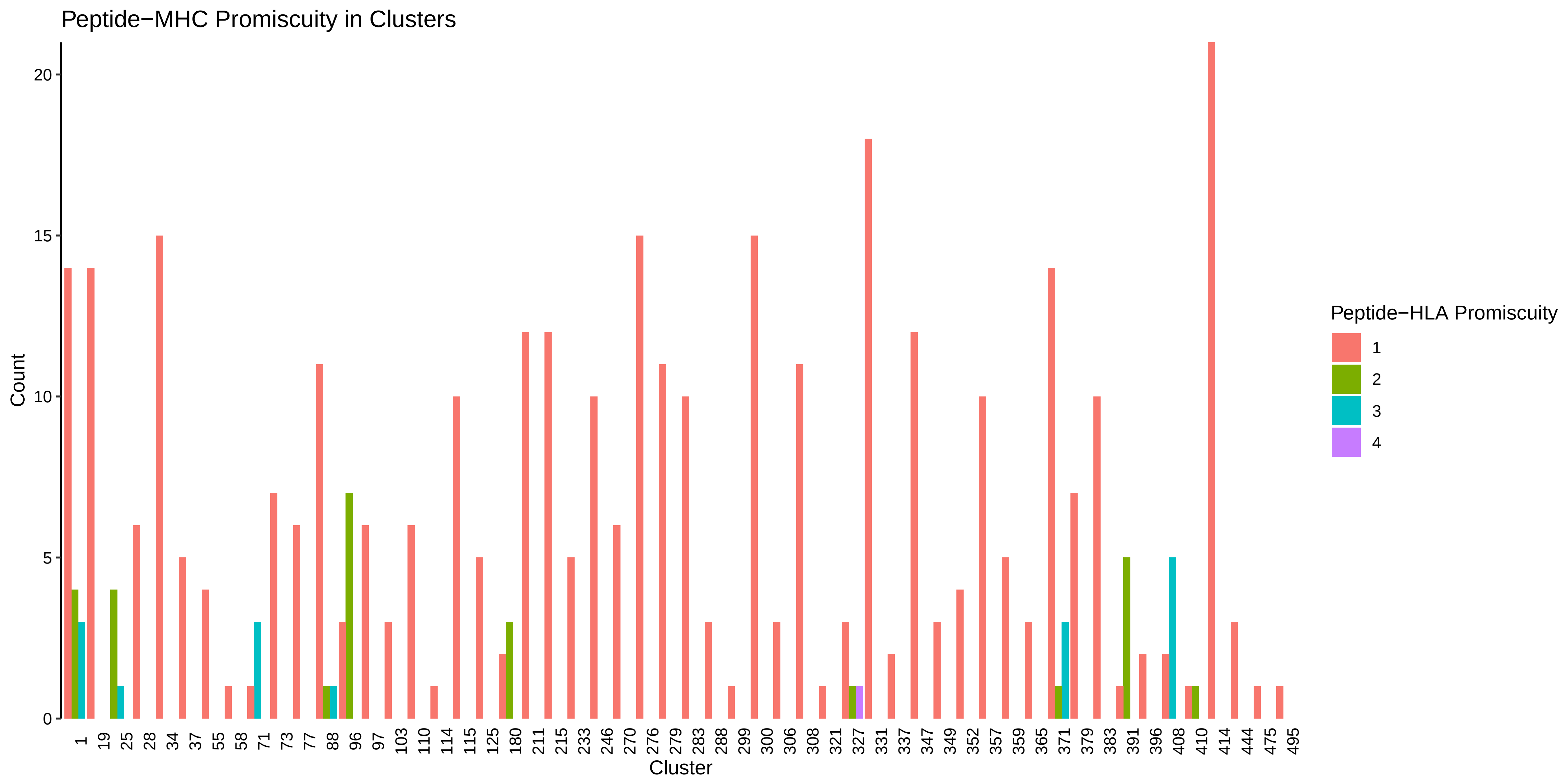

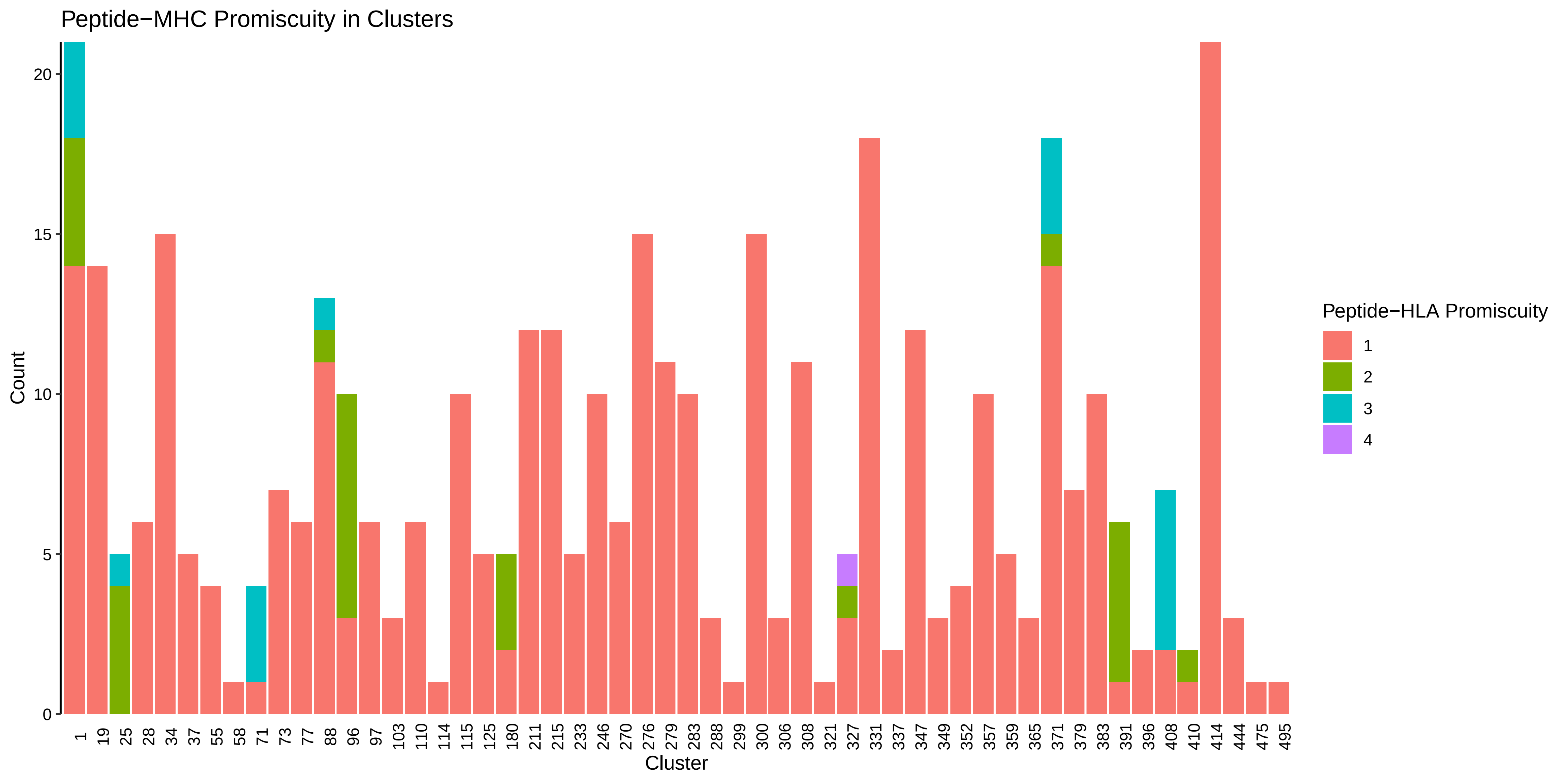

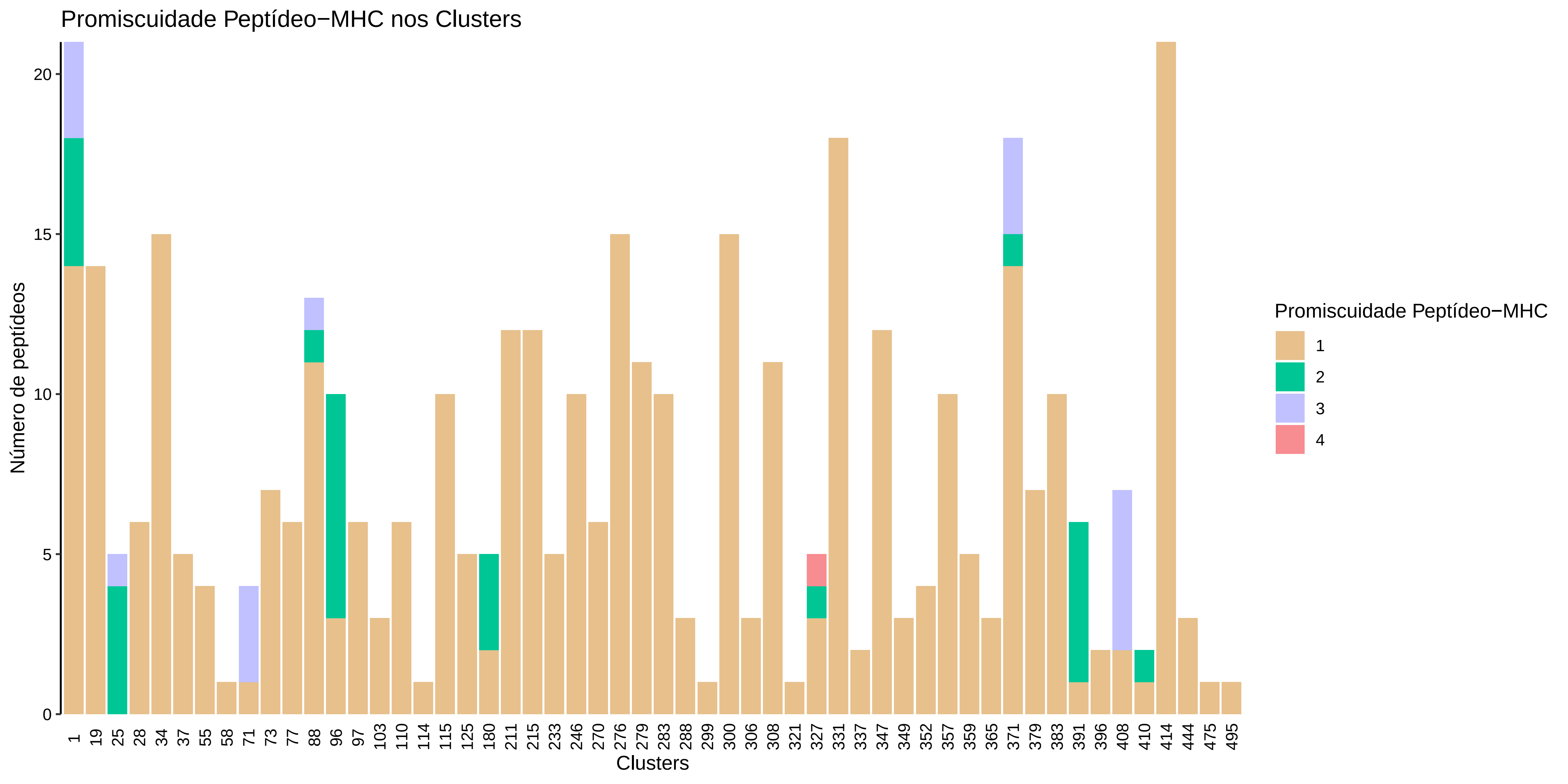
