## Supplementary Table I for "EpitoCore: mining conserved epitope vaccine candidates in the core proteome of multiple bacteria strains"

Supplementary Table I - Number of proteins predicted per step

| STRAIN | PROTEOME | TMHMM* | SHP | PSORT |
|---|---|---|---|---|
| TH135 | 4800 | 481 | 211 | 83 |
| OCU901s | 4569 | 496 | 192 | 84 |
| HP17 | 4561 | 490 | 197 | 87 |
| OCU873s | 4499 | 475 | 198 | 82 |
| OCU464 | 4754 | 481 | 191 | 81 |
| H87 | 4969 | 509 | 203 | 83 |
| MAC109 | 4841 | 497 | 209 | 87 |

*Predicted transmembrane proteins after SHP removal
